# Construction of whole genomes from scaffolds using single cell strand-seq data

**DOI:** 10.1101/271510

**Authors:** Mark Hills, Ester Falconer, Kieran O’Neil, Ashley D. Sanders, Kerstin Howe, Victor Guryev, Peter M. Lansdorp

**Affiliations:** Terry Fox Laboratory, BC Cancer Agency, Vancouver, BC V5Z 1L3, Canada; StemCell Technologies, Vancouver, BC V6A 1B6, Canada; AbCellera Biologics, Vancouver, BC, V6T 1Z4, Canada; Department of Pathology and Laboratory Medicine, University of British Columbia, Vancouver, BC, V6T 2B5, Canada; EMBL, Heidelberg, Germany; Genome Reference Informatics, Wellcome Trust Sanger Institute, Hinxton, Cambridge CB10 1SA, UK; European Research Institute for the Biology of Ageing, University of Groningen, University Medical Center Groningen, 9713 AV Groningen, The Netherlands; Department of Medical Genetics, University of British Columbia, Vancouver, BC, V6T 1Z4, Canada

## Abstract

Accurate reference genome sequences provide the foundation for modern molecular biology and genomics as the interpretation of sequence data to study evolution, gene expression and epigenetics depends heavily on the quality of the genome assembly used for its alignment. Correctly organising sequenced fragments such as contigs and scaffolds in relation to each other is a critical and often challenging step in the construction of robust genome references. We previously identified misoriented regions in the mouse and human reference assemblies using Strand-seq, a single cell sequencing technique that preserves DNA directionality^1, 2^. Here we demonstrate the ability of Strand-seq to build and correct full-length chromosomes, by identifying which scaffolds belong to the same chromosome and determining their correct order and orientation, without the need for overlapping sequences. We demonstrate that Strand-seq exquisitely maps assembly fragments into large related groups and chromosome-sized clusters without using new assembly data. Using template strand inheritance as a bi-allelic marker, we employ genetic mapping principles to cluster scaffolds that are derived from the same chromosome and order them within the chromosome based solely on directionality of DNA strand inheritance. We prove the utility of our approach by generating improved genome assemblies for several model organisms including the ferret, pig, Xenopus, zebrafish, Tasmanian devil and the Guinea pig.

---

The mouse^3^ and human^4^ genome references have revolutionized biomedical research, and facilitated many advances in studies of transcription, epigenetics, genetic variation, evolution and cancer^5^. However, while both assemblies are of very high quality, they still contain fragments that have not been localized to specific chromosomes, and large regions (typically flanked by unbridged gaps) that are incorrectly oriented with respect to adjacent scaffolds^1, 6^. These features highlight the difficulty in finishing genome maps, with typically repetitive or degenerate regions preventing robust overlapping/contiguous sequence across the length of the chromosome. As methods improve, assemblies themselves evolve over time as sequences are added, gaps are closed and errors resolved. For example, in the 13 years from the first public release of the complete human genome sequence (NCBI33)^7^ to the current assembly (GRCh38), the total number of represented nucleotides has only increased 2.79 % (82.27 Mb). While the change in genomic content between these two builds appears relatively modest, the change in the organization of the sequence has been dramatic. Regions with unknown local order and orientation have been corrected and placed, and incorrectly merged artefacts such as pseudo-duplications, misorientations and chimeras have been repaired. Correctly arranging available sequence data is therefore as important as uncovering new sequences in the process of improving genome references. Indeed, much of the drive to discover additional sequences revolves around the need to physically connect and orient contigs and scaffolds within the assembly, which is especially challenging within tracks of repetitive DNA. The methods involved in gap resolution and reorientation typically involve deeper sequencing of genomic DNA or BAC libraries^8^, but often also rely on novel methods such as optical mapping^9, 10^ and long-read sequencing technologies^11 12, 13, 14^. Recent studies have shown that improvements to optical mapping (termed whole-genome mapping) can facilitate *de novo* genome assemblies when used in conjunction with massively parallel sequencing^9^. This method involves creating scaffolds from sequencing libraries of genomic DNA and fosmid clones, followed by whole-genome mapping to match sequence patterns between contigs, generating super-scaffolds. While whole-genome mapping reduces the misorientation errors and can place scaffolds over a relative large distance, it is still mainly used as a verification tool, rather than the primary line of evidence used to produce chromosome-level genome references. With the increased availability and affordability of massively parallel sequencing (MPS) technologies, there have been efforts to build *de novo* assemblies from short read data. Ancillary methods to validate and expand these assembles are becoming increasingly important in this endeavor, as many of MPS assemblies show a marked reduction in quality and are dependent on the type of aligners used^15^. Given the relatively short sequence identity available to build contigs from MPS data, any nucleotide ambiguities can impact the alignment and affect the resulting assembly. Therefore, methods to detect incorrectly aligned scaffolds, to aid in creating the assembly and to provide secondary verification of the assembly are important to improve these strategies. Long read approaches resolve some of the ambiguity in joining overlapping reads into contigs^16^, but suffer from a higher nucleotide error rate that can mask overlapping regions between contiguous sequence, and still only cover a local region rather than the whole chromosome.

The single cell next-generation sequencing technique Strand-seq, offers an attractive orthogonal tool to refine and correct reference assemblies^1, 2^. Strand-seq involves sequencing parental DNA template strands in single daughter cells and the method preserves the directionality of DNA. This is achieved by culturing cells in the presence of BrdU, a thymidine analogue that is incorporated exclusively into newly formed DNA strands. After cell division, single cell libraries are created and treated with a combination of Hoechst and UV to remove the newly-formed strands, resulting in single-stranded library fragments containing template DNA only^17^. As replication is semiconservative, the DNA template strands that are inherited into daughter cells are either the Watson (W, ‘-’ or 3′-5′) or the Crick (C, ‘+’ or 5′-3′) strand^18^. By maintaining this directionality, we previously showed that Strand-seq locates sister chromatid exchanges (SCEs) at unparalleled resolution, seen as a template strand switching from W to C or vice versa^1, 17, 19, 20^. In addition, Strand-seq has been shown to have many applications including the mapping of polymorphic inversions^2^, haplotyping^21, 22^ and studies of DNA repair in yeast^23^ and humans^20^. These applications as well as the principle of genome assembly using Strand-seq data are illustrated in Figure 1. For the latter, the orientation of sequence reads is used to generate scaffolds, with sequence reads in each scaffold having either a WW, WC or CC state in every cell that is sequenced (Figure 1D)^24^. Similarly, any changes in strand state within a scaffold either represents an SCE event or an error where contigs have been incorrectly fused. When a strand state switch occurs at the same location in all libraries it can be delineated as an error, while an SCE event will occur randomly. SCE events are important elements in creating Strand-seq assemblies, as every scaffold downstream of an event will have a different state to everything upstream of an event (Supplementary Figure 1). Similar to meiotic recombination in genetic mapping approaches, this feature allows ordering of scaffolds along chromosomes.

**Figure 1:**
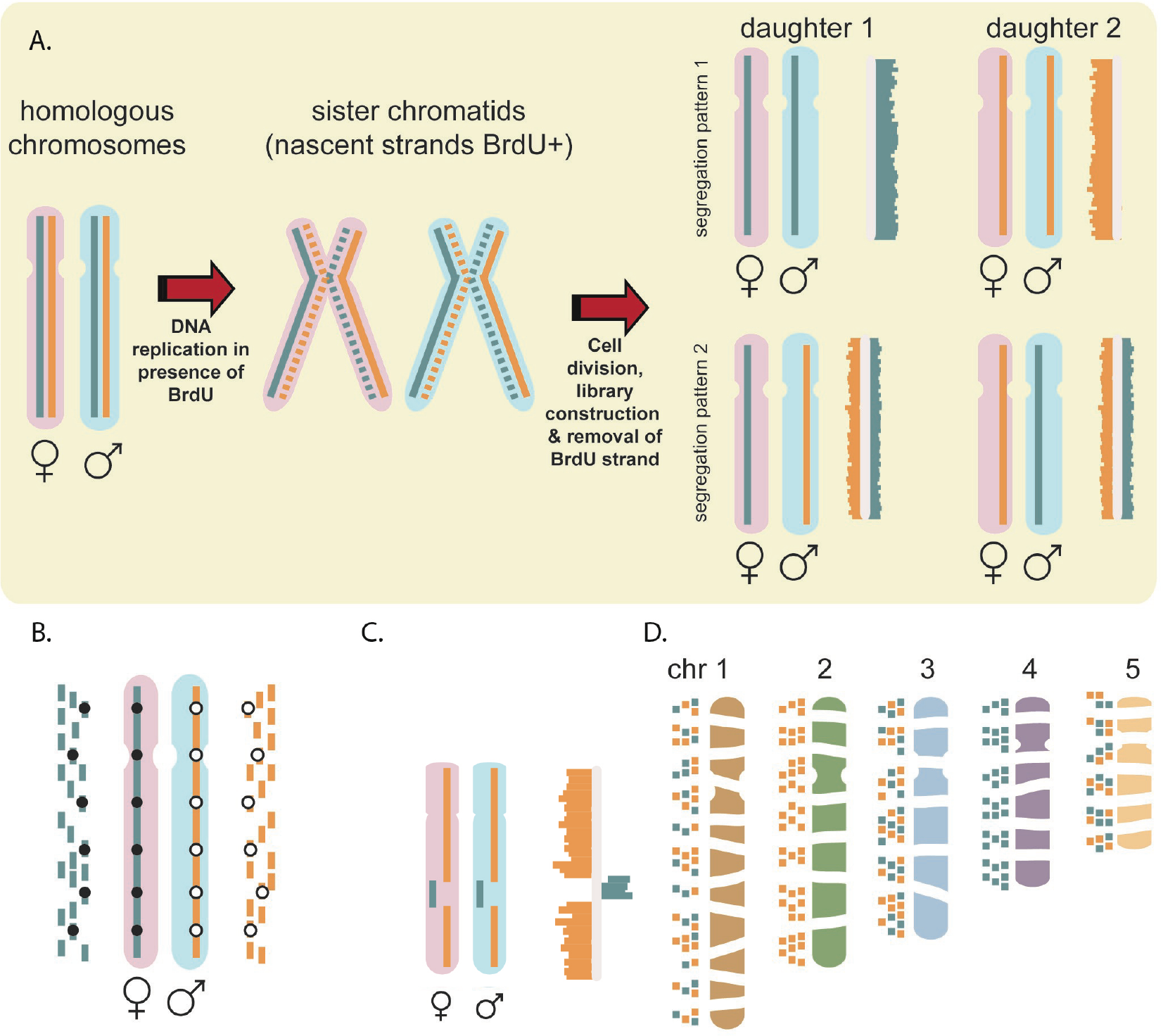
The principal and applications of Strand-seq. A. Strand-seq involves sequencing template strands. Parental homologues (pink & blue) are double stranded; Crick (C) strand in blue, Watson (W) strand in orange. DNA replication occurs in the presence of BrdU, which incorporates into the replicated strand (dotted lines). Sequencing libraries from single daughter cells have BrdU-containing strand selectively removed to generate directional chromosomes; either CC, WW (top) or WC (bottom) depending on segregation. Histograms of directional reads are plotted on ideograms for each chromosome. B. When homologues inherit different template strands, haplotypes can be determined. In the example, all C reads map to the maternal homologue so all SNVs identified (black dots) form the maternal haplotype, and all W reads map to the paternal homologue, so all SNVs identified (white dots) form the paternal haplotype. C. Structural variation can be identified in Strand-seq libraries. Inversions will align to the opposite strand of the reference assembly as so be identified as a change in template strand state D. Strand-seq can be used to create assemblies since contigs from the same chromosome will have the same template inheritance pattern. Grouping based on shared template inheritance patterns determines which fragments belong together. Note in the example contigs from ch1, chr3 and chr5 have the same template pattern (WC) so require additional libraries to establish which contigs belong to which chromosome.

Previously, Strand-seq was used to resolve orientation errors in the GRCm37 assembly to which the data were aligned^1^. In addition, we were able to map many of the remaining unlocalized and unplaced scaffolds from this assembly by matching the template inheritance pattern of the fragments to the inheritance pattern of individual chromosomes^6^. Supporting data verified the presence of the misorientations identified by Strand-seq^1^, and the Mouse Genome Reference Consortium incorporated this information into subsequent builds. For the human genome, orienting fragments in the reference assembly is complicated by common polymorphic inversions^2^. Nevertheless, using Strand-seq, we identified 41 reference assembly misorientations and/or minor alleles (allele frequency < 0.05) in GRCh37, which were distinguished from > 100 polymorphic inversions found in unrelated individuals^2^. Strand-seq was also used to assemble haplotypes along the entire length of all chromosomes without generational information or statistical inference^21, 22^. While we have utilized Strand-seq to correct polished assemblies, it is more complicated to align scaffolds together in the absence of a whole assembly map. However, our ability to successfully improve near complete assemblies motivated us to apply Strand-seq to other species with less complete, draft-quality genome builds.

Many organisms that are important for biomedical research have very incomplete genome assemblies. Here, we have applied Strand-seq and the bioinformatics analysis package contiBAIT^24^ to aid in refining the assemblies for six such organisms (Table 1). To demonstrate the ability of Strand-seq to generate robust assemblies by clustering thousands of unconnected contigs, three organisms were selected with scaffold-stage assemblies at different levels of completeness. The ferret (*M. putorius furo*) assembly consists of 7,783 unplaced scaffolds^25^ and is an important model for studies of human respiratory diseases, including influenza infection and transmission. The assembly of the Tasmanian Devil (*S. harrissii*) genome has been spearheaded to aid in studies of an atypical transmissible cancer, Devil Facial Tumour Disease, which is decimating the population. Currently this assembly contains 35,974 scaffolds placed to chromosomes, but without a specific order^26^. Finally, the Guinea pig (*C. pocellus)*, is an important model organism used in the study of vaccines and the research and diagnosis of infectious diseases. This assembly consists of 3,142 large unplaced scaffolds^27^.

**Table 1.**
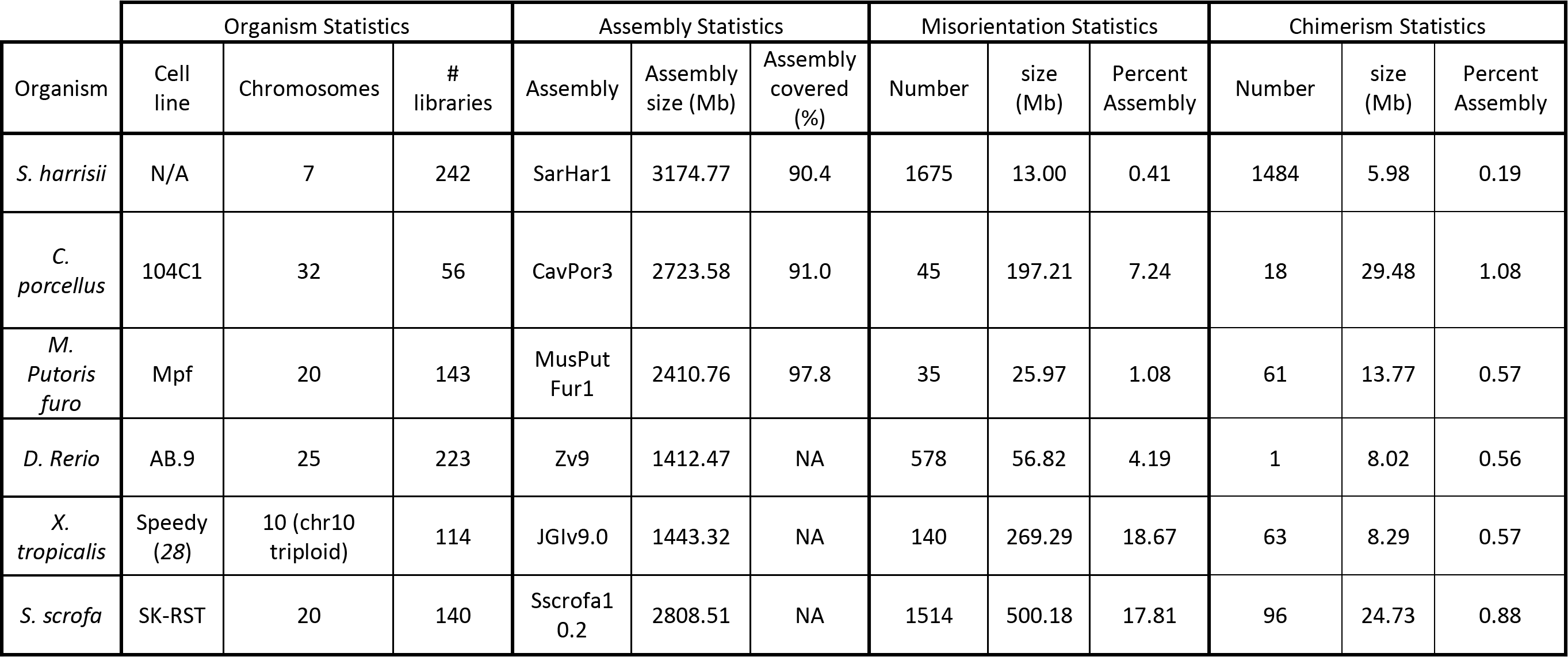
Summary of data from all six organisms. The organism statistics outline the cell line used, the number of Strand-seq libraries used in the study and the expected number of chromosomes. The chromosome number was adjusted based on the expected allosomes for the gender of the cell line for each organism. The assembly statistics includes the assembly that the Strand-seq libraries were aligned to, the (gapped) size of that assembly, and the proportion of scaffolds covered in the data (where applicable). The misorientation and chimera statistics highlight the number, genomic size and proportion of the assembly affected by misorientations and chimeric fragments respectively.

We further used Strand-seq to correct misorientations and incorrectly placed scaffolds in three chromosome-stage assemblies. The principle of this approach is based on arranging scaffolds into linkage groups (Figure 2) and ordering them along the full length of each chromosome (Supplementary Figure 1). Of these six organisms we used for enhancing genome references, the pig (*S. scrofa*) was selected for its significance in agriculture and in medicine, as well as in understanding evolution during animal domestication. Most of the sequence (92 %, 5,344 scaffolds), has been ordered into the 20 chromosomes, with a further 4,562 scaffolds remaining unplaced. However, this assembly still contains 69,541 spanned and 5,323 unspanned gaps^28^. Since there is no underlying information on the orientation of scaffolds separated by unspanned gaps (which have no supporting evidence for the orientation of the contigs they flank), this would suggest that as least some of the scaffolds are incorrectly oriented. Genome references of many other important model organisms also built on the chromosome-level contain multiple gaps and unplaced fragments. For example, the zebrafish (*D. rerio*) is an important model in vertebrate development and gene function, and while the zebrafish assembly^29^ (Zv9) is of high quality and mostly complete, it included 1,107 unplaced fragments (55.4 Mb), and 3,427 unspanned gaps. A further model organism with a large research community, Xenopus (*X. tropicalis*), has an assembly with more unplaced fragments (6,811, 167.9 Mb), but no unspanned gaps^30^.

**Figure 2:**
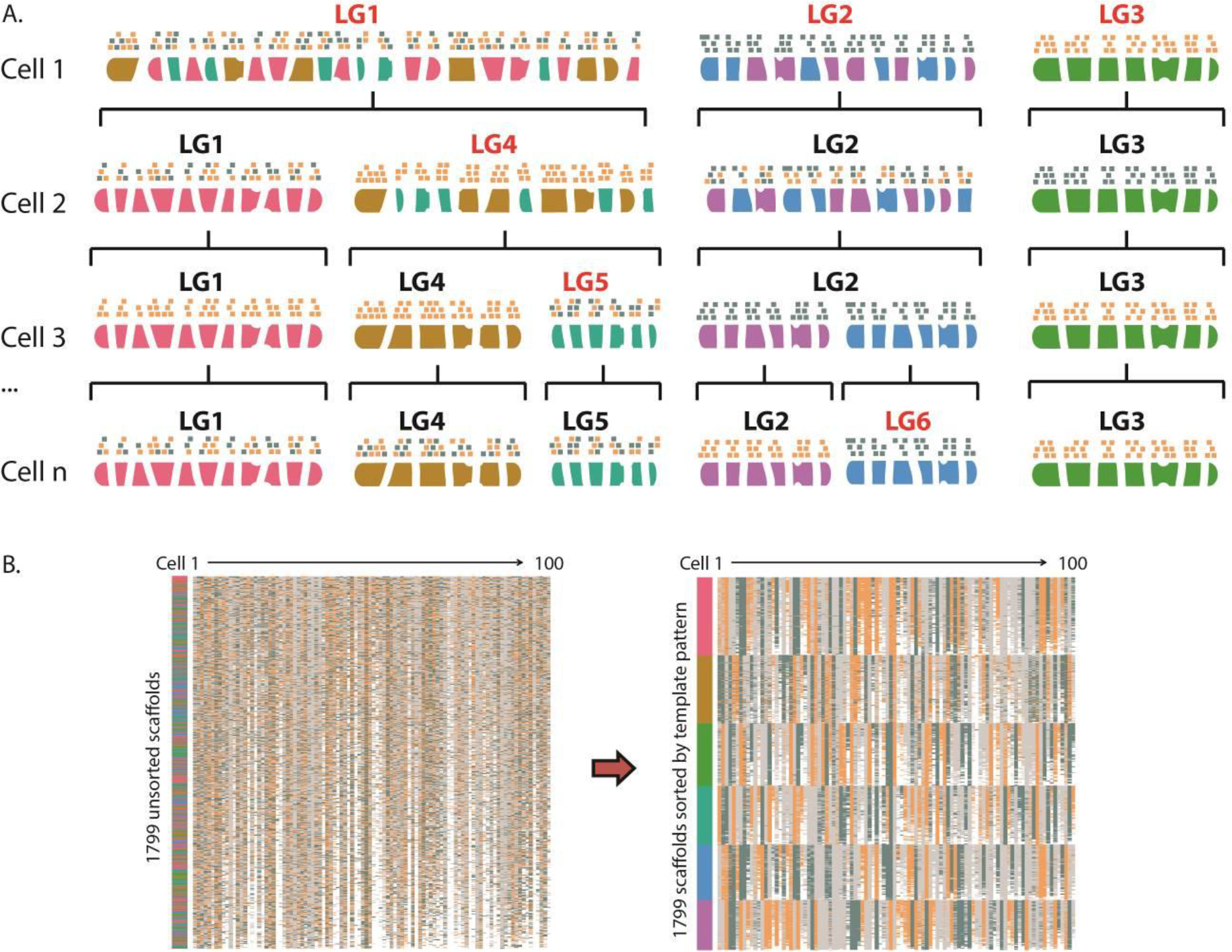
Clustering scaffolds based on Strand inheritance. A. Schematic for clustering 6 chromosomes. Each chromosome pair will harbor one of three template inheritance states: WW, WC or CC (W=blue, C=orange). Through analysis of the template inheritance pattern of multiple cells, scaffolds from the same chromosome share the same pattern and can be resolved. For example, in Cell 1, three chromosomes are represented in LG1, but are resolved in subsequent cells. B. Subsetted data showing 1,799 unsorted ferret scaffolds belonging to six linkage groups across 100 cells (CC=blue, WW=orange, WC=grey, no data=white). Prior to clustering (left plot), scaffolds from the same chromosome are unknown, while after clustering (right plot), scaffolds that share template inheritance patterns across individual cells are resolved. Vertical color bar represents called members for each of the six linkage groups.

## Results

For the six organisms studied, we built and used Strand-seq libraries from between 56 to 242 single cells per species (Table 1), and data were aligned to their respective assemblies and analyzed using the Bioconductor package contiBAIT^24^ (Table 1). We also included previously published mouse^1^ and human^2^ Strand-seq datasets as positive controls. For all organisms, we were able to correct multiple errors that encompassed large regions of these assemblies (both between and within scaffolds). We achieved this by identifying two distinctive signatures that represent common errors that propagate within assemblies (Figure 3). First, regions that showed consistent and complete reversal in template state for a portion of the scaffold were flagged as a misorientation (or as an polymorphic inversion between the cell line sequenced and the assembly). Next, regions that showed no inheritance similarity with neighbouring sequence were identified as putative chimeras that arise from contig mis-joins such that portions of scaffolds are placed to the wrong chromosome. For the former, misoriented sequences were reoriented within the fragment and flagged as errors in the assembly (Figure 3). For the latter, chimeras were split at the mis-join site and independently clustered to identify the correct location of these fragments (an example chimeric scaffold is shown in Supplementary Figure 2).

**Figure 3:**
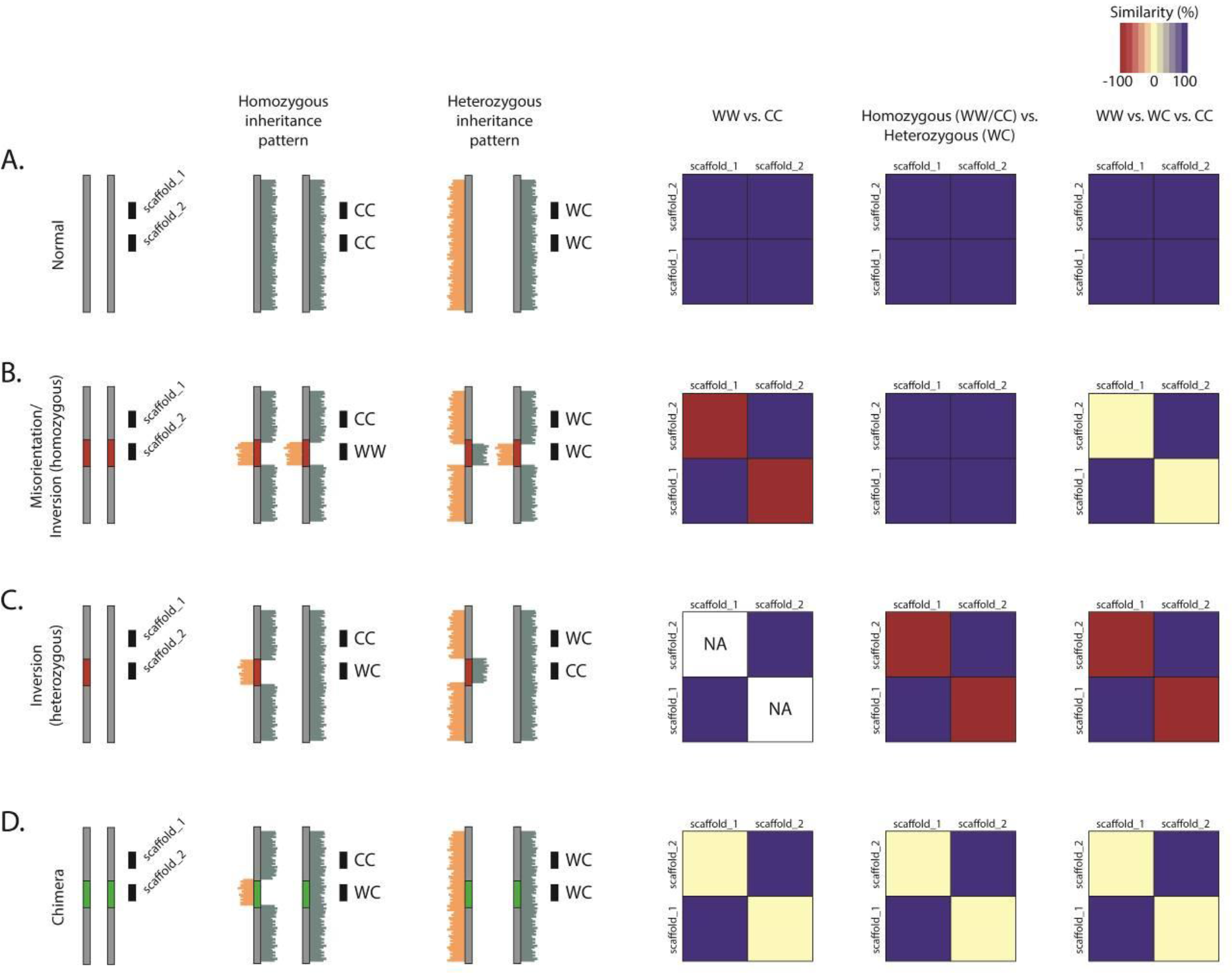
The effect of different assembly errors or structural variation on clustering. Different errors will generate characteristic patterns in the clustering data. Consider two scaffolds in close proximity on a chromosome, scaffold_1 and scaffold_2. A. In a case where both scaffolds are oriented in the same direction, the scaffolds will have the same strand-state patterns. When comparing homozygous patterns (WW scaffolds against CC scaffolds), heterozygous patterns (WW or CC scaffolds against WC scaffolds) or comparing all three strand states against each other, there will be high similarity. B. In the case of a misorientation (or a homozygous inversion), the strand-state patterns will be antithetical when comparing homozygous states, as whenever scaffold_1 is WW, scaffold_2 will be CC, and as such, these scaffolds will be completely dissimilar. However, since misorientations are not visualized in heterozygous inheritance patterns, when comparing WW or CC states against WC states, the scaffolds are highly similar. When comparing all three states against each other, the similarity seen with WC scaffolds and dissimilarity seen with WW or CC scaffolds will cancel out, resulting in ~50% similarity. C. In cases of a heterozygous inversion, either scaffold_1 or scaffold_2 may have a homozygous state, but not both. Therefore, no comparisons can be made when only considering the homozygous states, and NA values are generated. There will however, be a high degree of dissimilarity when comparing homozygous and heterozygous states. It is important to distinguish these natural structural variants from assembly reference errors. D. In cases where a scaffold is incorrectly located to a chromosome (i.e. a chimera), the inheritance pattern between the two scaffolds will be random, and there will be no significant similarity or dissimilarity between these scaffolds.

Using the template inheritance as a bi-allelic marker for every scaffold in the respective assemblies, we devised a method to cluster scaffolds based on the expectation that those belonging to the same chromosome will show the same bi-allelic template pattern across multiple Strand-seq libraries^24^. To achieve this, all fragments from a single Strand-seq cell were divided into one of three groups based on the inheritance patterns of their templates: WW, CC, or WC, and then grouped and ordered based on shared inheritance states between all fragments and across all cells (Figure 2). In this way, we were able to assign each scaffold to a linkage group (LG), where all scaffolds within the same LG belonged to the same physical chromosome. The software is able to account for the fact that assembly scaffolds may be in 5′-3′ or 3′-5′ orientation and reorients fragments into the same directions. These LGs are therefore equivalent to a ‘super scaffold’: they encompass many scaffolds and fragments that cluster together, are oriented in the same direction and represent a draft chromosome (Figure 2). Moreover, since the strand inheritance pattern is a feature of the entire chromosome, Strand-seq is able to resolve scaffold associations along entire chromosomes rather than at a megabase level. For each scaffold assembly, the majority (> 90%) of fragments clustered together into the same number of LGs as there are chromosomes from that organism (Figure 1). For example, for the ferret genome (20,XX), 97.9 % of the assembly fragments mapped to the 20 largest LGs (Figure 4, Supplementary Figure 3). Each of these 20 groups represent scaffolds that have been correctly oriented and show co-inherited strand states, consistent with them belonging on the same chromosome. Similarly, 90.9 % of Guinea Pig (32,XX) scaffolds mapped to the 32 largest LGs (Supplementary Figures 3 & 4), and 90.4 % of Tasmanian Devil (7,XY) assembly fragments mapped to 7 LGs (Supplementary Figure 5). Since unlocalized and unplaced fragments are not tethered to whole chromosome scaffolds, the orientation of these fragments was expected to be mostly random. Our data supported this, showing there were approximately equal numbers of unlocalized and unplaced fragments represented in each direction (Figure 5B). Using the same methodology, we were able to locate many of the unlocalized fragments present in the chromosome-stage assemblies for the pig, zebrafish and Xenopus.

**Figure 4:**
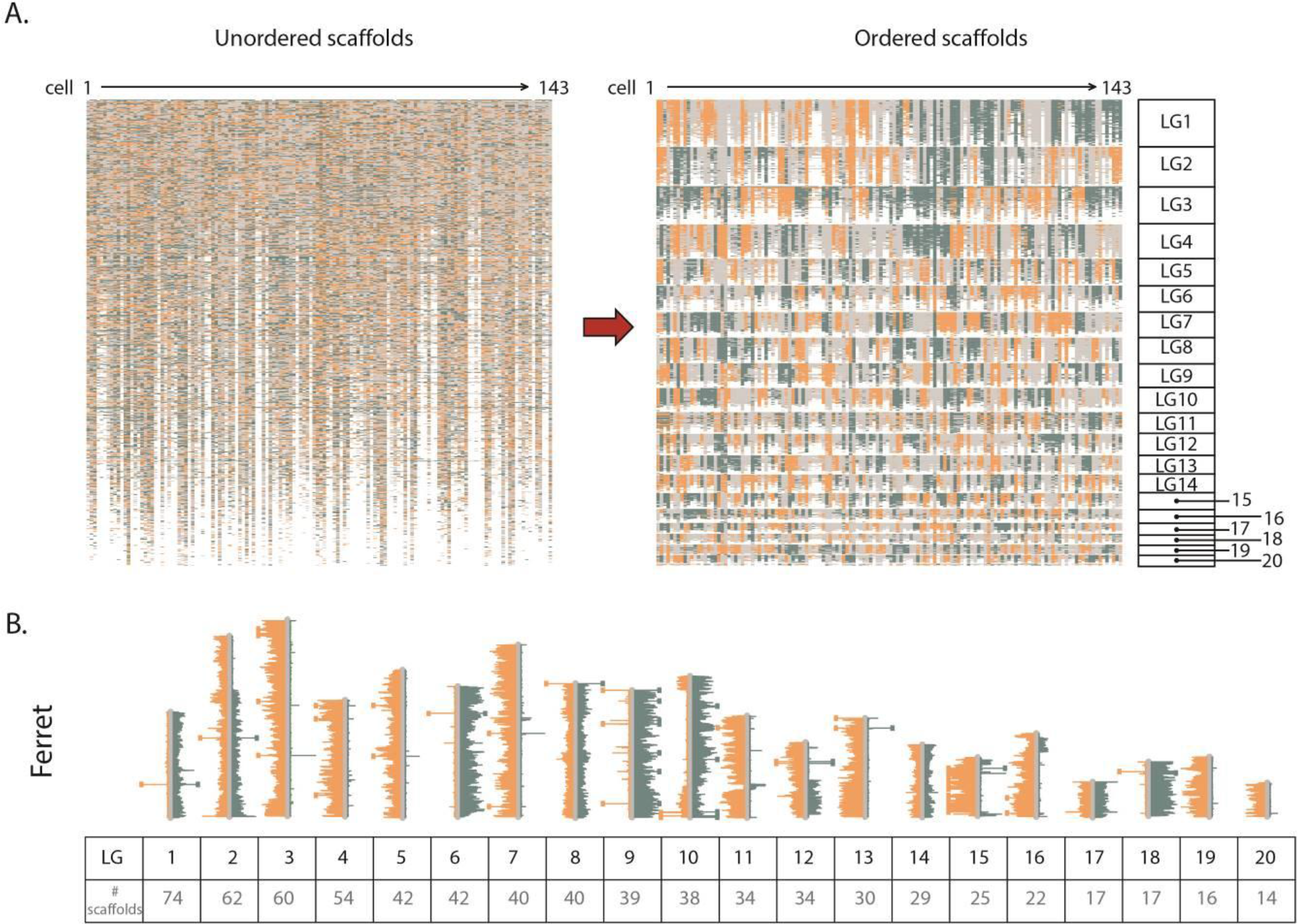
Assemblies made from non-contiguous scaffolds based on Strand-seq data. A. Left panel shows ferret scaffolds presented in the current assembly order. Orange, blue and grey represent scaffolds with WW, CC and WC reads respectively. Right panel shows scaffolds after contiBAIT reordering B. Representative ideogram plot of a ferret library after clustering and ordering scaffolds. Each linkage group is represented by a certain number of scaffolds. Chromosomes with WW, WC and CC inheritance patterns are observed in this library. Changes in strand state represent SCE events and are used to map the relative locations of scaffolds.

Misorientations were identified in all assemblies, though to varying degrees (Figure 5, Table 1). By conventional methodologies, orienting contiguous sequences flanked by gaps has been difficult, with BAC end sequencing being the primary approach to bridging these gaps. It was therefore not unexpected that the majority of misoriented scaffolds we identified occurred between assembly gaps. However, misorients were also identified within contiguous sequences, albeit at a lower rate. For example, we discovered 578 misoriented regions in the zebrafish assembly Zv9 (56.8 Mb, 4.19 % of the assembly), but only 22 of these were not flanked by gaps. To investigate our ability to correctly orient scaffolds using Strand-seq, we performed BioNano optical mapping and shotgun sequencing on a separate zebrafish cell line and compared scaffolding calls. More than 97 % of misorientations identified by Strand-seq were cross validated by at least one orthologous technique. Of these, 240 (41 %) were identified in shotgun sequenced clones, and 256 (44 %) were identified through BioNano optical mapping. Based on these data, our Strand-seq results were included as a validation method, and the misorientations identified were incorporated into the GCRz10 build of this genome.

**Figure 5:**
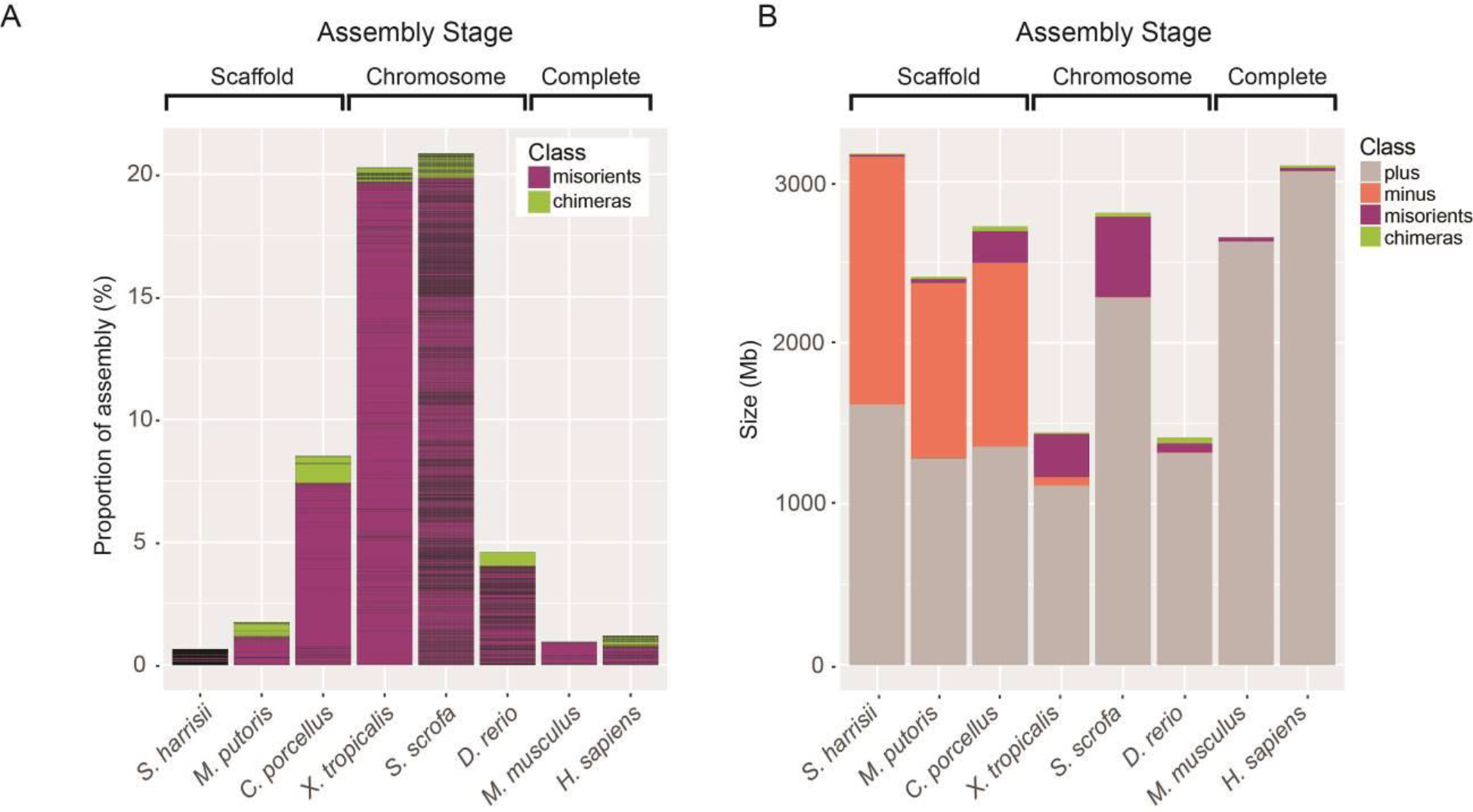
Assembly misorientations and chimeras are prevalent in early-stage genomes. A. Percentage of assembly fragments classified as misorients or chimeras. Horizontal lines represent the sizes of each error within the assembly. Note that all chromosome-level assemblies displayed multiple orientation errors. The chimeric fragment within zebrafish is derived from an inverted region in the AB strain with respect to the Tübingen assembly^33^, while misorients in the mouse were identified previously^1^, and chimeras and misorients identified in the human sample correlated with previously identified heterozygous and homozygous inversions respectively^2^. B. Barplot of scaffold orientation within each assembly. The predominant orientation of scaffolds within the assembly is set as correct (“+ strand”, grey), and the frequency of scaffolds that do not match this orientation is calculated. Misorients are subdivided into entire scaffolds that are in the opposite orientation to the majority of assembly scaffolds (dark green), and fragments within contiguous sequence that are in the incorrect orientation (purple). Chimeric fragments (green) are defined as portions of contiguous sequence that display a different template strand inheritance pattern and are therefore likely placed to an incorrect chromosome. The proportion of incorrectly oriented scaffolds constitute half of the scaffold-level assemblies. Chromosome-and complete-level assemblies have fewer scaffolds (higher N50 values), so most assembly errors occur within contiguous sequences.

Similar observations were made with the other chromosome assemblies: the pig reference (Sscrofa10.2) exhibited a greater degree of misorientation than the zebrafish assembly, with 1,514 fragments (500.18 Mb, 17.81 %) identified within the chromosome scaffolds. In addition, 96 chimeric fragments were discovered (24.73 Mb), split and relocalized. For the Xenopus assembly, 140 misorientations were found (269.29 Mb, 18.67 %) and 63 regions were flagged as chimeric. Using these data, we generated refined versions for each assembly, and after realigning, all Strand-seq reads were in the correct direction (Supplementary Figure 6). The quality of the scaffold-stage assemblies studied varied markedly based on misorientation and chimerism analysis (Figure 5, Table 1). For the Guinea pig reference, 18 putative chimeras were detected, while 45 misorientations (197.21 Mb, 7.24 %) within the scaffolds were found. Fewer misorientations were seen in the ferret assembly, with 35 identified (25.97 Mb, 1.08 %), while 61 chimeras were detected. Finally, we identified 1,675 putative misorientations in the Tasmanian devil assembly (13.0 Mb, 0.41 %) and a further 1,484 putative chimeras (Table 1).

As a final application of Strand-seq, we were able to organise scaffolds into a relative order within LGs. Using SCEs that naturally arise in single libraries and occur randomly during replication^1^, the template strand similarity between scaffolds from multiple libraries will progressively diminish the further apart they are in physical distance, as the likelihood of SCEs occurring between them increases. In this way, our approach is similar to classical linkage mapping, where genetic distance can be inferred as a function of the number of SCEs between two fragments (Supplementary Figure 1). Since chromosomal locations of all scaffolds had already been determined, we ordered these fragments based on SCE within each chromosome (Supplementary Figure 1). All data are included as bed files, which encompass the distinct LGs and order of fragments for each scaffold assembly, along with the directionality of all fragments for both scaffold and chromosome-level assemblies (Supplementary Data File).

## Discussion

The quality of genome assemblies is determined by the methods employed to build them, the algorithms used to create contigs and chromosomes, and the complexity of the genome. Genomes with high levels of repetitive elements have the potential to be assembled erroneously resulting in fused chimeric contigs, and genomes with segmental duplications can be collapsed or overrepresented as multiple copies^31^. Algorithms used to build contigs from overlapping sequences can vary wildly^15^, often resulting in chimeric contigs which may be retained in future builds.

Our results show that the quality of each original assembly is highly variable, which likely derives from the complexity of the genome, the type of technologies used for sequencing/scaffolding, and the algorithms used to build the assemblies^32^. Moreover, while model organisms often have homogeneous genomes due to inbreeding, the genetic heterogeneity of outbred organisms complicate and confound assembly strategies. Nucleotide variation can interfere with the joining of contigs, but more drastically, large polymorphic structural variation can impede the ability to create a reliable assembly (Figure 3). This kind of structural variation is prevalent within the human population, where 1.2 % of the genome (34.91 Mb) represents regions in which polymorphic inversions have been detected^2^. It is possible that sequencing a variety of outbred animals and creating a composite assembly will therefore result in conflicting scaffold joins, with inter-animal structural variation confusing the orientation and location of fragments. As such, the hybrid approach used for the pig assembly may explain the large degree of misorientations we observed. Here, the data, primarily derived from a female Duroc sow, were combined with sequence from four other porcine breeds; Large White, Meishan, Yorkshire and Landrace^28^. The AB zebrafish cell line used in our study was from a different strain than was used for the Tübingen assembly^33^, and we identified a previously described^33^ polymorphic peri-centromeric chromosomal inversion on chromosome 3 (chr3:46,945,080-56,227,809, data not shown). While we are unable to exclude the possibility that the assembly misorientations identified in our study are homozygous polymorphic inversions, other methods including *de novo* assembly through sequencing are also not immune to this issue. Furthermore, while heterozygous inversions can resemble contig mis-joins, they can be resolved since they display unique patterns within our data (Supplementary Figure 6). By combining this approach with Strand-seq haplotyping^21, 22^, we will be able to further resolve and phase these structures during the assembly process, although an initial assembly with which to align to is still necessary.

Using Strand-seq we have developed a novel approach to building assemblies that is completely independent of overlapping contigs. This approach can rapidly locate and localize fragments with as little as a single lane of a sequencing run. The ability to improve reference assemblies using common sequencing platforms is an advantage of Strand-seq over orthologous methods that require specialized equipment such as long-read sequencing methods and optical mapping. Furthermore, these results highlight that Strand-seq can assess contiguous sequences using multiple reads spread across fragments, and as such can readily identify incorrect contig misjoins. This approach has more in common with traditional genetic mapping strategies than standard assembly approaches, and can be applied to assemblies at the contig, scaffold, chromosome or complete stages. By identifying the order of scaffolds, this method will further aid in efforts to sequence across gaps using targeted PCR-based or long-read strategies. Collectively, we show that this approach simultaneously stratifies, orients and corrects assemblies. As the field relies more and more on computational assembly building from shorter massively parallel sequence reads, the opportunity for incorrect dovetail joining of overlaps to introduce chimeric contigs is increased. Taken together, our results show that Strand-seq is an effective approach for improving genome assemblies by allowing, in combination with other sequencing methods, to immediately correct, orient and link fragments together.

## Data Availability

Raw data from this study is available at EBI ArrayExpress (https://www.ebi.ac.uk/arrayexpress) under following accession: E-MTAB-6480.

## Author contributions

M.H conceived and designed the study. E.F, A.D.S. and P.M.L helped with data generation and study design. K.O. helped with design of bioinformatics approaches and data analysis. K.H. performed optical mapping studies in zebrafish and V.G. helped with data analysis. The manuscript was written by M.H. with assistance from all authors. P.M.L. supervised the project.

## Acknowledgements

Financial support for this work was provided by an Advanced Grant from the European Research Council to P.M.L.

---

**Supplementary Figure 1:**
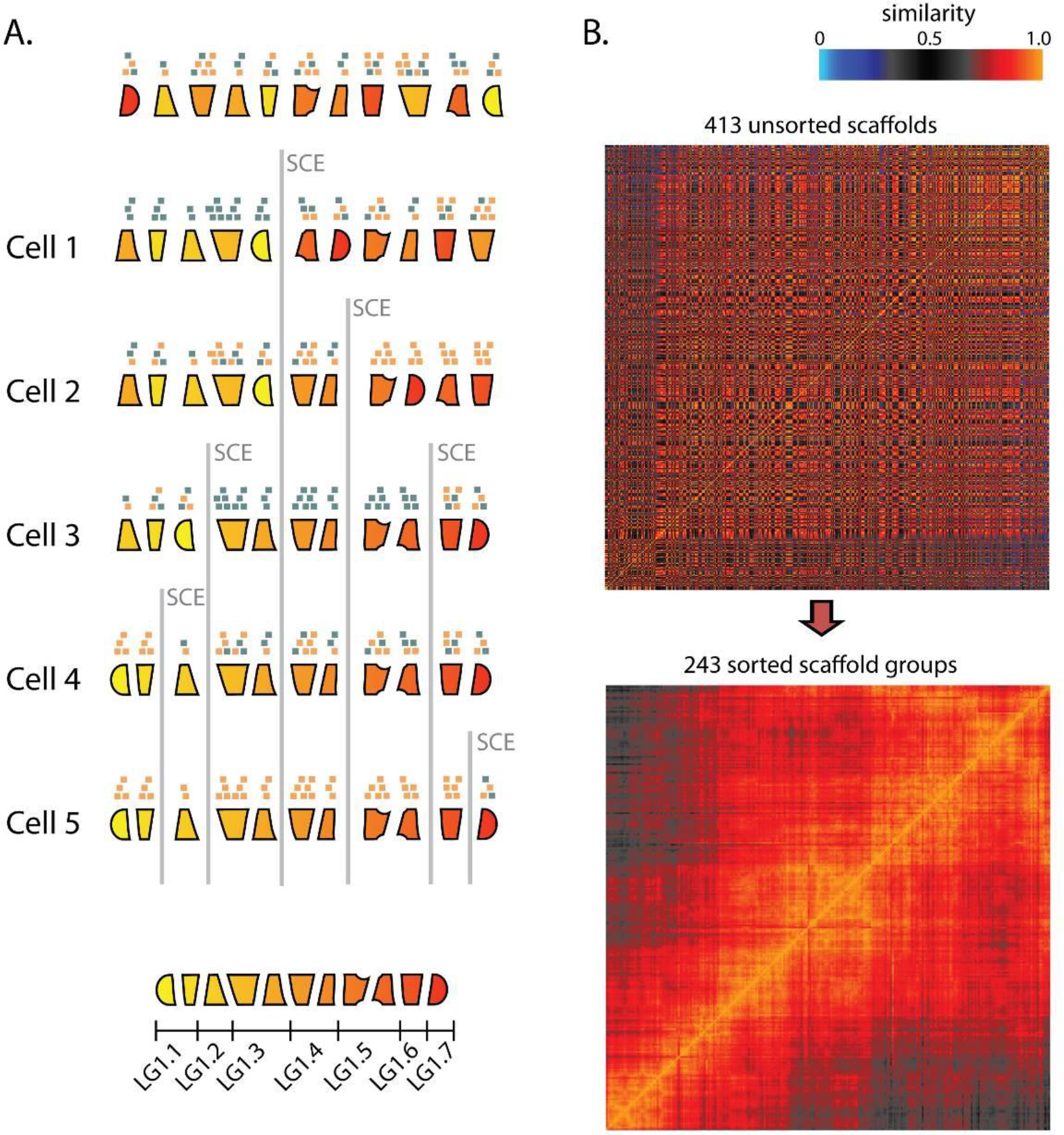
Clustering scaffolds within Linkage Groups. A. Schematic for clustering 11 scaffolds within Linkage Groups (LG). Across 5 different cells, SCEs reshuffle the pattern of template strands between scaffolds. Scaffolds are clustered into groups of similarity upstream and downstream of an SCE event. As more SCEs are included, more ordered subgroups are resolved. Clustering is performed through a Monte Carlo algorithm. B. Example heat plots of ferret LG consisting of 413 scaffolds. While some scaffolds show identical template patterns (due to their close proximity), 243 scaffold groups can be elucidated and ordered. The greatest degree of template strand similarity is between nearby scaffolds, with progressively weakening linkage between more distant scaffolds.

**Supplementary Figure 2:**
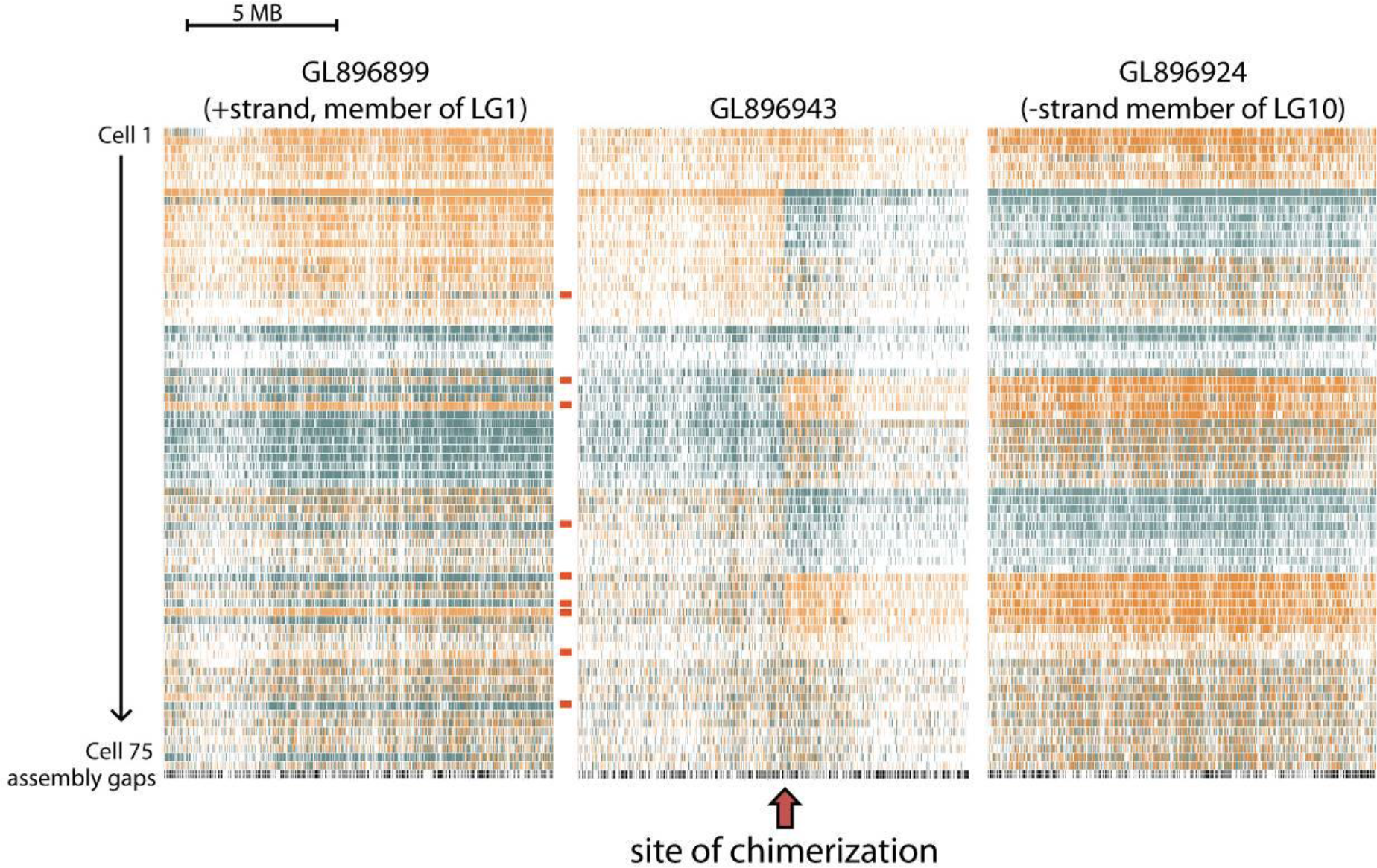
Example of chimeric scaffold in the ferret. GL896943, a ~12 MB scaffold in the ferret assembly, was flagged as a putative chimera due to a change in template strand state occurring at the same location in multiple cells (red arrow). Our software splits these scaffolds at the site of chimerization into two smaller contigs prior to clustering. In this example, one contig matched scaffolds belonging to LG1 (GL896899 used as a representative member), and the other matched scaffolds belonging to LG10 (GL896924 used as a representative member). Note the nine differences (red bars) between GL896899 and GL896943, occurred from SCEs between these fragments that are distantly located within the LG.

**Supplementary Figure 3:**
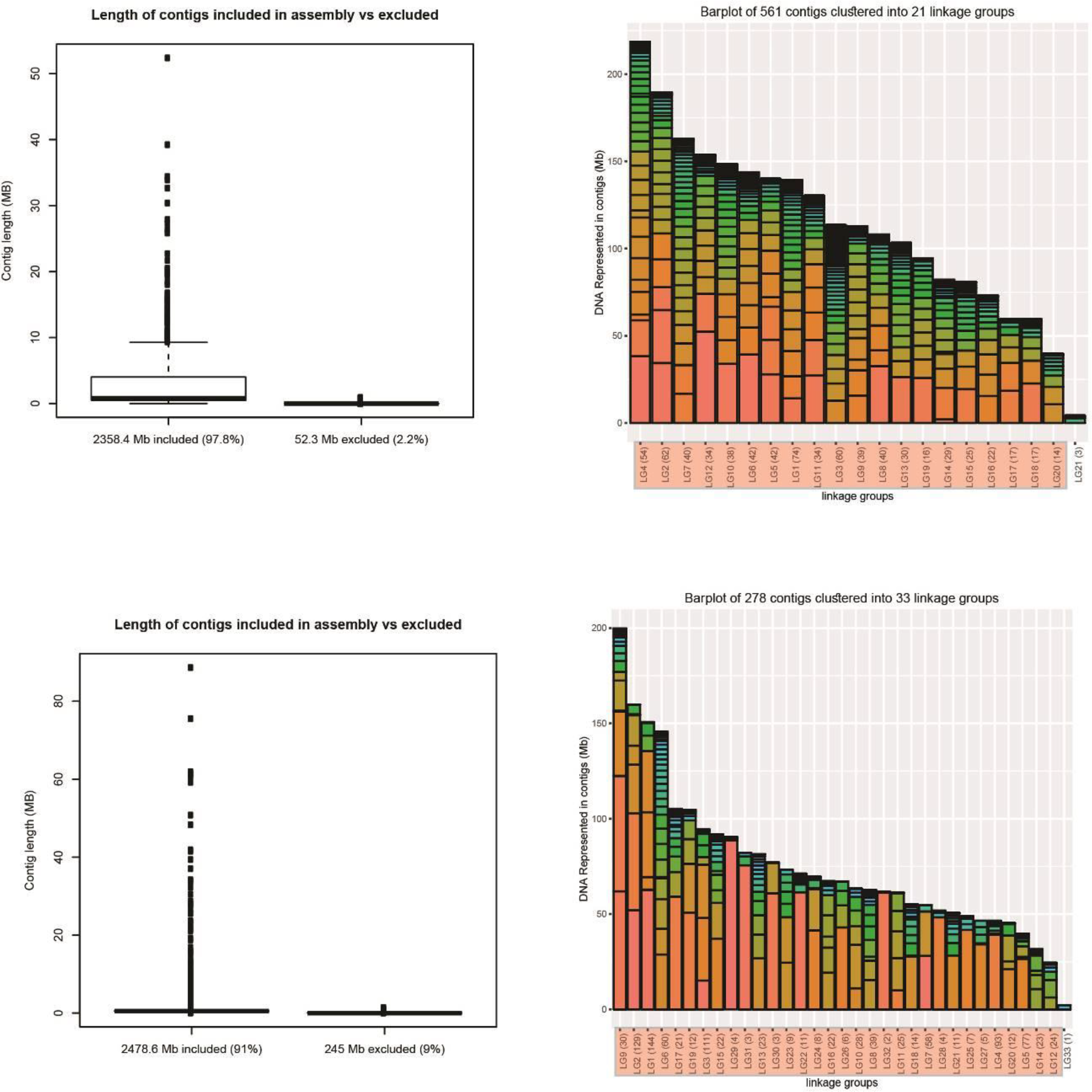
Scaffolds clustered into chromosome-sized linkage groups (LGs). The ferret (top) and Guinea pig (bottom) assemblies consist of thousands of scaffolds of no known location. Boxplots show that the majority of scaffolds are placed into LGs by Strand-seq analysis, with generally small fragments excluded (colors used to distinguish scaffold sizes). Bar plots show the distribution of scaffolds that have clustered together into LGs based on template inheritance patterns. LGs highlighted in red represent the expected number of chromosomes for that organism.

**Supplementary Figure 4:**
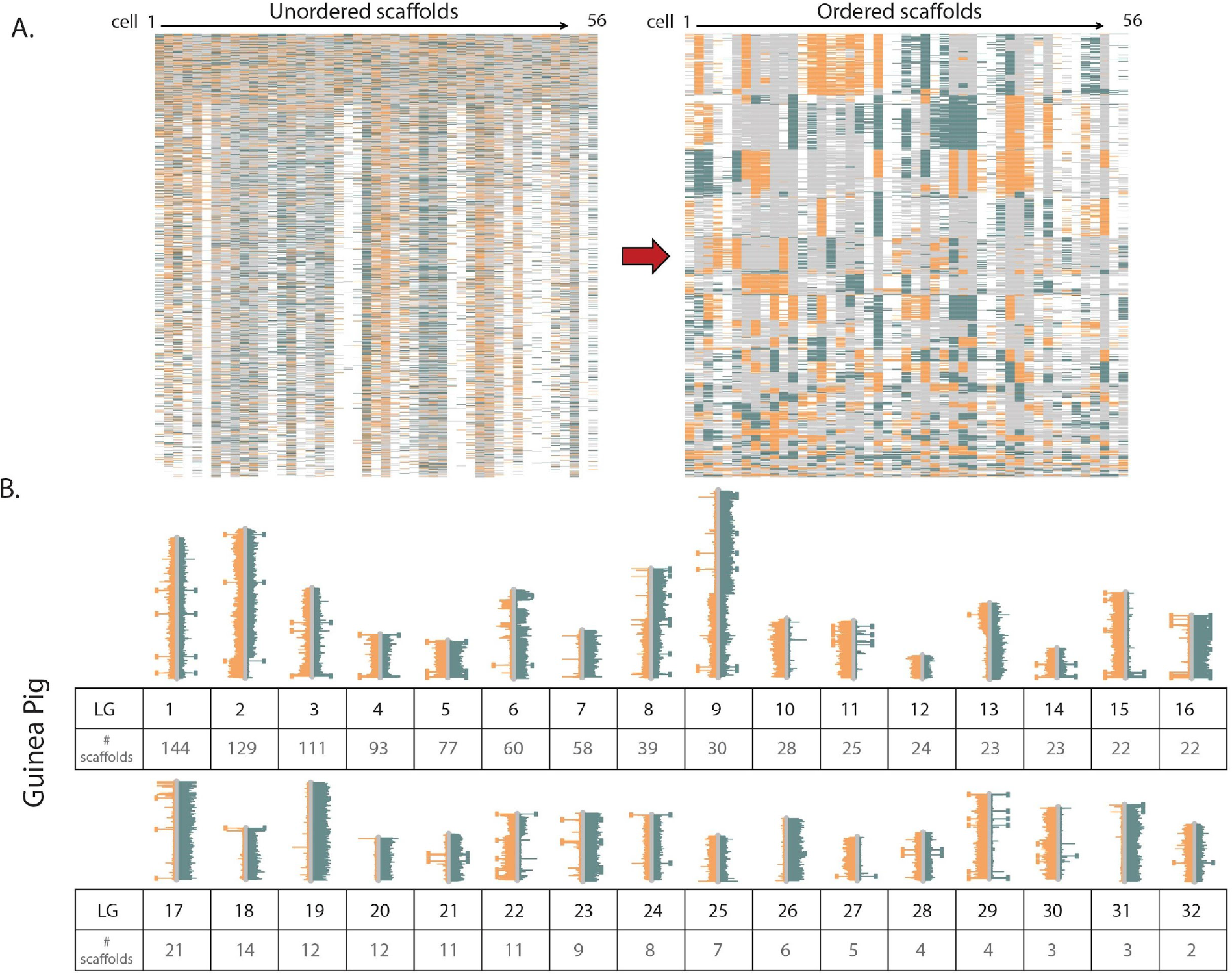
Guinea pig assembly made from non-contiguous scaffolds based on Strand-seq data. A. Left panel shows Guinea pig scaffolds presented in the current assembly order. Right panel shows scaffolds after contiBAIT reordering B. Representative ideogram plot of a Guinea pig library after clustering and ordering scaffolds. Each linkage group is represented by a certain number of scaffolds. Chromosomes with WW, WC and CC inheritance patterns are observed in this library. Changes in strand state represent SCE events.

**Supplementary Figure 5:**
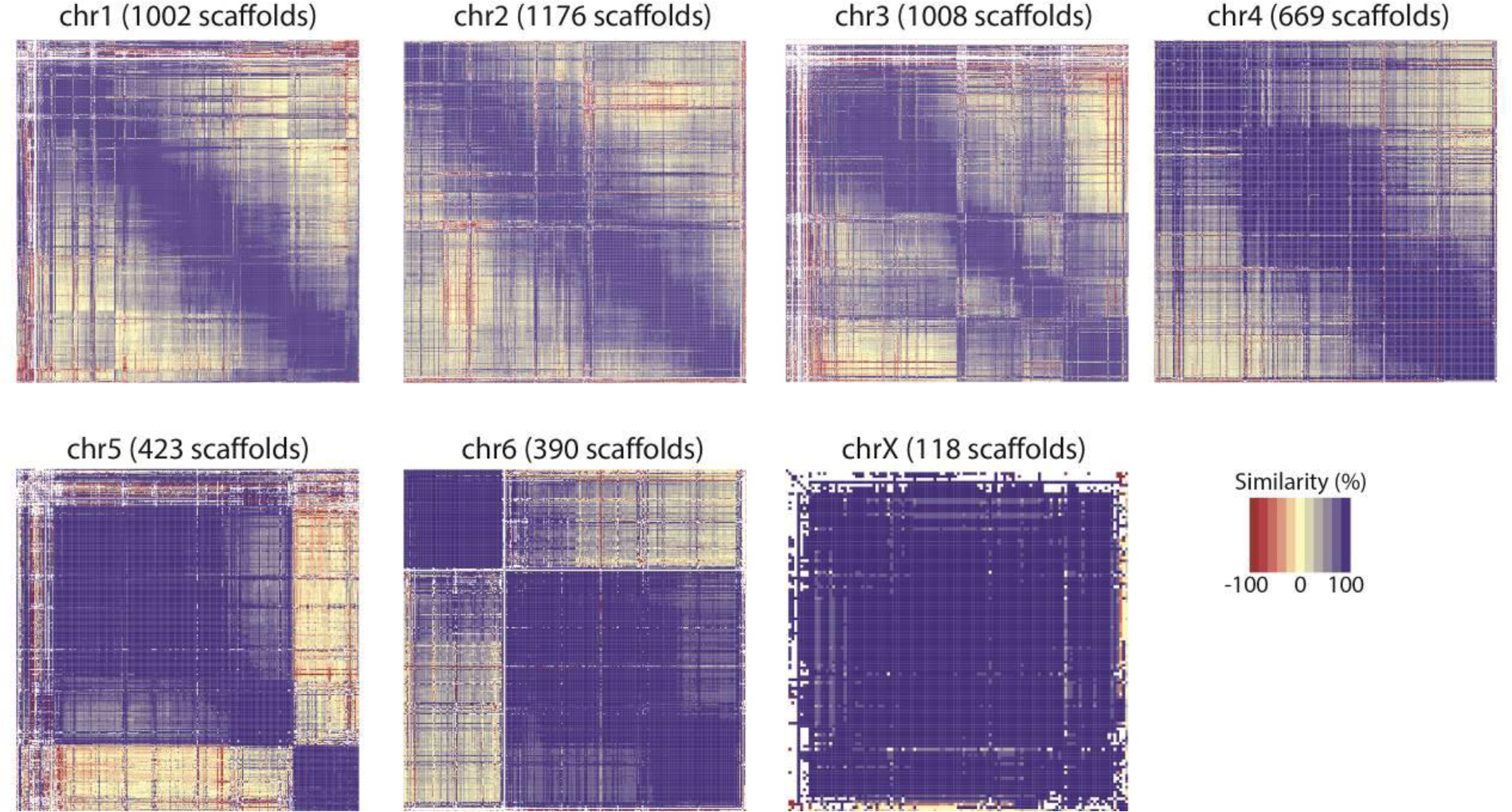
Tasmanian devil scaffolds ordered within chromosomes. While the Tasmanian devil assembly consists of 35,974 scaffolds, most of these fragments have been located to specific chromosomes within the assembly (6 autosomes and an X chromosome). Heat maps show the relative order of these fragments within this assembly. A total of 4,786 scaffolds were clustered, which represents 90.4 % (2,869.1 Mb) of the Tasmanian Devil assembly.

**Supplementary Figure 6:**
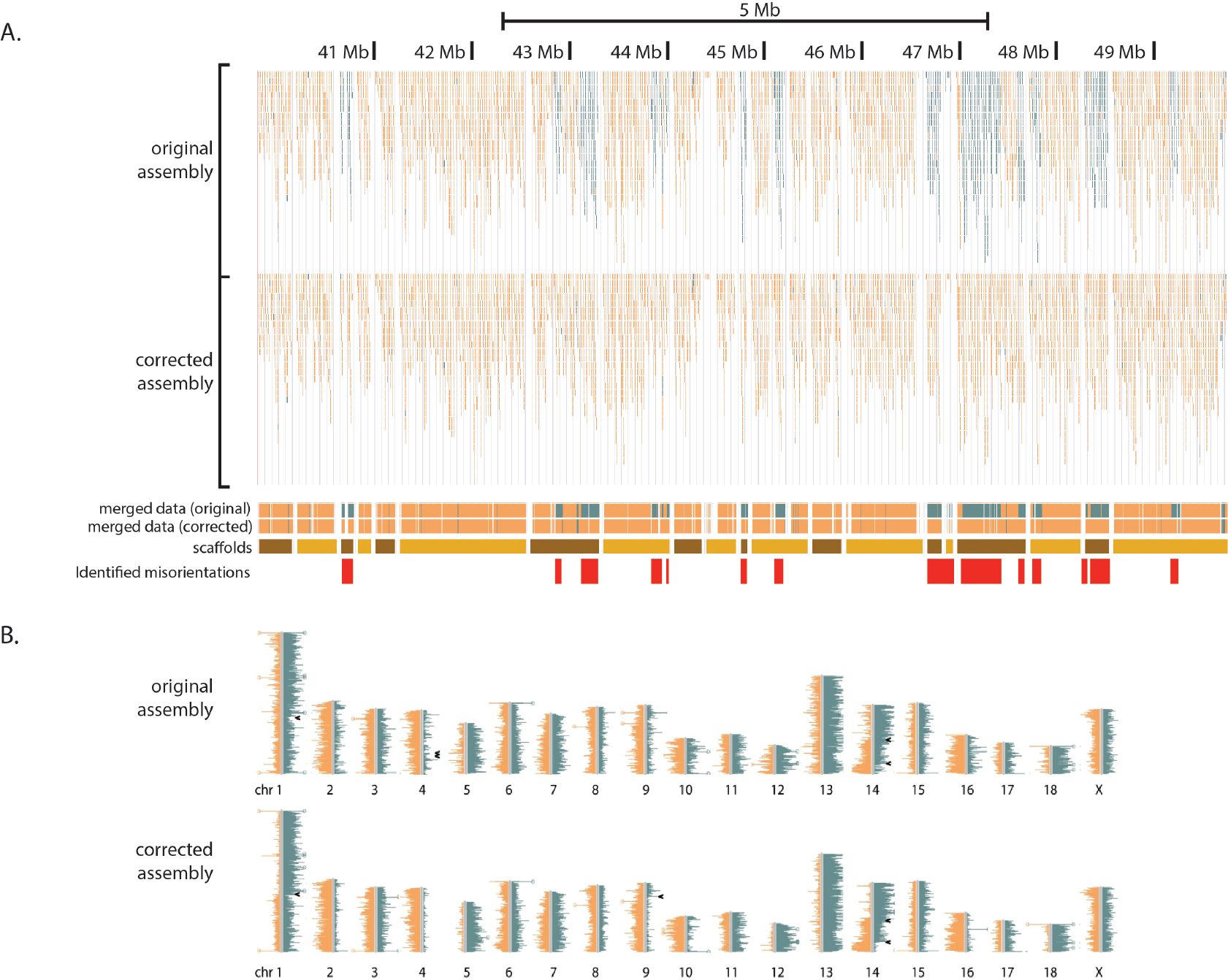
Correcting the orientation of scaffolds in the pig assembly. A. 17.81 % of the pig assembly was identified as misoriented. By reverse complementing regions on misorientation, we have corrected this assembly with Strand-seq data. BED plots show read distribution and directionality from a UCSC genome browser screenshot. Red boxes represent locations identified at misorients. Note these events occupy both entire scaffolds, or portions of scaffolds (suggesting errors within contiguous sequence). B. BAIT ideograms for a single cell generated before and after correction of the pig assembly. The template strand inheritance pattern cannot be distinguished in the original assembly due to high prevalence of WC states, but is resolved after reorientation. Arrowheads indicate locations of sister chromatid exchange events.

